# Geographical and ecological drivers of coexistence dynamics in squamate reptiles

**DOI:** 10.1101/2022.01.27.478006

**Authors:** Laura R. V. Alencar, Tiago B. Quental

**Affiliations:** Departamento de Ecologia, Instituto de Biociências, Universidade de São Paulo, São Paulo, SP, 05508-900, Brazil; Department of Ecology & Evolutionary Biology, Yale University, New Haven, CT, 06511, USA

**Keywords:** dispersal ability, competition, niche divergence, snakes, lizards, islands, continents

## Abstract

**Aim:** Species richness varies widely across space. To understand the processes behind these striking patterns, we must know what are the relevant drivers underlying species coexistence. Several factors can potentially shape species coexistence such as the speciation process, the time since divergence between lineages, environmental effects, and intrinsic properties of the organisms. For the first time, we model the coexistence dynamics of lizards and snakes across broad temporal and spatial scales, investigating the role of species interactions, dispersal ability, and geographic area.

**Location:** Global

**Time period:** Last 20 million years

**Major taxa studied:** Squamata (lizards and snakes)

**Methods:** We used 448 closely related species pairs and their age since divergence across 100 dated phylogenies. We categorized each pair as sympatric or allopatric and as occurring on islands or continents. We measured morphological traits to quantify niche divergence and used range and body size as proxies for dispersal ability. We applied a model-comparison framework in lizards and snakes separately to evaluate which factors best explained their coexistence dynamics.

**Results:** We found that distinct factors drive the coexistence dynamics in lizards and snakes. In snakes, species pairs that coexist tend to occur on islands and are more different in body size, suggesting that both geographical setting and species interactions might be relevant factors. In contrast, we only found evidence that dispersal ability shaped the coexistence of lizards, where species coexist when they have higher dispersal abilities.

**Main conclusions:** Lizards and snakes greatly differ in coexistence dynamics. Higher heterogeneity in coexistence dynamics within lizards and group-specific life-history aspects might help to explain these findings. Our results emphasize that the interaction between where organisms are and who they are, ultimately shapes biodiversity patterns. We also highlight interesting avenues for further studies on species coexistence in deep time.

## Introduction

Biodiversity is heterogeneously distributed across the Earth, and to explain why certain regions comprise more species than others has been one of the major challenges in Ecology and Biogeography (Gaston, 2000; Ricklefs, 2004). At global and regional scales, the number of species is determined by speciation, extinction and migration (Wiens & Donoghue, 2004; Ricklefs, 2006). However, to fully understand such spatial variation in species richness we also need to know the mechanisms that ultimately promote species coexistence and generate the observed species richness patterns at finer spatial scales (Weber & Strauss, 2016; Pigot et al., 2018). How and why the distribution of species changes across space and time has been the focus of intense debate in the literature (e.g. Jackson & Overpeck, 2000; Sexton et al., 2009; Louthan et al., 2015).

Several factors can potentially shape the distribution of organisms and consequently shape species coexistence (Hutchinson, 1957; Vamosi et al., 2009; Wisz et al., 2013). Distinct speciation modes, for example, are linked to distinct predictions regarding the patterns of coexistence between two closely related species. Sympatric speciation assumes that two species will coexist since their emergence (Grossenbacher et al., 2014), whereas allopatric speciation will produce geographically isolated species (Kozak & Wiens, 2006) that can eventually expand their distributions and subsequently coexist (Pigot & Tobias, 2015; Weber & Strauss, 2016; Pigot et al., 2018). In this case, the time since divergence between lineages can play an important role in determining coexistence patterns.

Biotic interactions and the intrinsic properties of organisms, such as their dispersal ability, can also affect the process of geographic expansion, either accelerating or preventing species coexistence over time (Johnson & Stinchcombe, 2007; Vamosi et al., 2009; Lowe & McPeek, 2014; Jønsson et al., 2016; Weber & Strauss, 2016). From an ecological perspective, the importance of competition in shaping coexistence is based on the “principle of competitive exclusion”, which suggests that species occupying similar ecological niches would not be able to coexist because they require the same limited resources to survive (Hutchinson, 1957; Hardin, 1960). The outcome of these ecological processes on an evolutionary time-scale could be character displacement among populations or a pair of species (Brown & Wilson, 1956; Schluter, 2000), species sorting where only species with different ecological niches will coexist (Anderson & Weir, 2021), or even the extinction of an entire lineage (e.g. Silvestro et al., 2015). Indeed, the number of studies suggesting that competition is a relevant factor in shaping the distribution of organisms at global and macroevolutionary scales has rapidly increased (Gotelli et al., 2010; Pigot & Tobias, 2013; Silvestro et al., 2015; Pigot et al., 2018). Pigot and Tobias (2013) for example, found that the coexistence of closely related species of birds seems to be mediated by differences in their ecological niche. On the other hand, other factors might play a role, and these same authors later found that dispersal ability also determine how fast two species might expand their distributions and coexist (Pigot & Tobias, 2015; Pigot et al., 2018). Dispersal is indeed widely suggested to play a central role in the redistribution of organisms, contributing to the colonization of new areas and range shifts, representing a potentially important component driving coexistence dynamics (Lowe & McPeek, 2014; Jønsson et al., 2016; Weber & Strauss, 2016).

Although biotic interactions and dispersal ability have been frequently invoked to explain both spatial and temporal biodiversity patterns, the geographical scenario where species thrive is another important aspect to be considered. Islands are widely known for producing unique biodiversity patterns given their smaller size and geographic isolation compared to continental settings (MacArthur & Wilson, 1967; Losos & Ricklefs, 2009; Baeckens & Van Damme, 2020). Changes in body size like gigantism and dwarfism are frequently reported in insular vertebrates (“island rule”, Foster, 1964; Van Valen, 1973; see also Benítez-López et al., 2021 but see Meiri, 2007). Patterns like these are suggested to be the result of several mechanisms characterizing insular systems, such as reduced predation, relaxed inter-specific competition and resource limitation (Losos & Ricklefs, 2009; Baeckens & Van Damme, 2020; Benítez-López et al., 2021). However, the consequences of these insular particularities on the coexistence dynamics of species are less clear (Ricklefs, 2010; Pigot et al., 2018). On one hand, it is feasible to expect that coexistence might be achieved faster on islands compared to continents given that the first comprise smaller geographical limits and is usually characterized by a relaxation of biotic constraints. On the other hand, islands harbor populations that might be more likely to suffer stochastic extinctions or introgression possibly erasing any signs of sympatry (Pigot et al., 2018).

Several other ecological, physiological, and behavioral aspects can also influence species coexistence dynamics (Gaston, 2000; Buckley et al., 2012). Squamate reptiles (i.e. lizards and snakes) are ectothermic animals and their distributions will frequently be limited by their physiological requirements (Buckley et al., 2012; Vitt & Caldwell, 2014). As a result, their distributions can be strongly determined by climatic conditions (e.g. Buckley et al., 2012). Lizards and snakes, however, comprise more than 10.000 species ranging across a wide spectrum of latitudes and biomes (Roll et al., 2017; Uetz & Hosek, 2017). While the species richness of snakes shows the typical latitudinal gradient similar to endothermic organisms, lizards show a pattern distinct from any other tetrapod group, with a higher richness in Australia (see Roll et al., 2017). Furthermore, although closely related, snakes and lizards differ greatly in their morphology and in several aspects of their natural history (Sites et al., 2011; Vitt & Caldwell, 2014). Generally speaking, snakes have a slower metabolism and lower population density compared to lizards (Vitt & Caldwell, 2014); lizards comprise more diverse diet habits, including insectivorous, carnivorous and even herbivorous species (Pianka & Vitt, 2003; Meiri, 2008), whereas snakes are strictly carnivorous (Greene, 1997). This impressive diversity is actually an indication that species distributions in squamates might not be regulated by the same processes, and that aspects not related to climate might also play an important role in shaping species distributions (e.g. Algar et al., 2013; Cunningham et al., 2015). *Anolis* lizards, for example, comprise a classic example of squamate reptiles among which species coexistence is strongly determined by interspecific competition (Losos, 2011).

To reveal the relevant factors shaping the coexistence dynamics of a diverse group such as squamates, and whether snakes and lizards share or not these factors, we need investigations across broader taxonomic and spatial contexts. Here we take this approach by investigating the role of biotic interactions, dispersal ability and different geographical settings in shaping the coexistence dynamics across closely related species of squamate reptiles. To do that, we use a model-comparison framework to (1) evaluate the main speciation mode (i.e sympatric or allopatric) in lizards and snakes; (2) test if the tendency to geographically overlap over time increases in species that are ecologically distinct, (3) have high dispersal abilities, and/or (4) occur on islands. Our study is the first to explicitly model the coexistence dynamics of squamates over time in broad temporal and spatial scales. On top of that, we take into account the possible effects of different ecological and geographical scenarios in driving coexistence. Our results provide insights into the factors shaping species distributions and the dynamics of coexistence, contributing to a better understanding of the processes shaping biological communities and, at a broader scale, of those that modulate the evolution of biodiversity.

## Methods

### Closely related species pairs

To explore coexistence dynamics in squamates, we used one of the most complete molecular phylogenies (Tonini et al., 2016) to define a pool of species pairs (“sister” species), representing the most closely related species in the maximum-likelihood topology. Our pool included only species from well sampled genera (70% or more of all described species of a given genus had to be in the phylogeny) and for which the phylogenetic placement of species pairs was highly supported (equal to or higher than 0.95). To calculate genus sampling, we followed the taxonomy of the Reptile Database up to January 2017 (Uetz et al., 2017). We were able to identify 538 species pairs (161 of snakes and 377 of lizards).

### Geographical ranges

To determine if species comprising a given species pair coexist, we used species polygon range maps mostly from Roll et al. (2017). For an additional 20 species, whose maps were not available in Roll et al. (2017), we used those provided by the IUCN Red List Assessment. We used the range map provided by Birskis-Barros et al. (2019) for the South American rattlesnake *Crotalus durissus*, given the outdated distribution for this species in the previous two databases. From the 538 species pairs, we could not obtain range maps for four pairs given that at least one of the species did not have a map available.

From the 534 species pairs for which we had range maps, we excluded species that occur in marine or mangrove environments (10 pairs) and terrestrial pairs where species are separated by marine barriers (species occurring on different landmasses or islands, see Pigot & Tobias, 2015) (those represent 57 pairs). The dispersal dynamics characterizing these species pairs might potentially differ from those where species are terrestrial and occur on the same landmass and would potentially add noise to our analyses.

### Age since divergence

To model coexistence over time, we used the age since divergence of each species pair estimated by Tonini et al. (2016) as our temporal measurement. Given that estimates of age since divergence can be highly uncertain, we randomly chose 100 different phylogenies from those generated by Tonini et al. (2016) to obtain a range of possible age estimates for our species pairs. It is important to mention that these 100 phylogenies were generated using the same backbone molecular phylogeny and, therefore, the relationships between species pairs remain the same, but the age estimates differ (see Tonini et al. 2016 for details). Therefore, we incorporated in our analyses 100 different age since divergence estimates for each species pair (see below).

To avoid including species pairs for which evolutionary history might have been strongly influenced by extinction we set a divergence time limit to our species pairs. We first took the median of the 100 ages since divergence estimates for each pair and kept those pairs comprising a median equal to or lower than 20 million years ago. After all curatorial work, our final pool comprised 448 pairs of species (132 of snakes and 316 of lizards) that we used to investigate the coexistence dynamics in squamates (see Appendix I). These 448 pairs span a wide diversity of taxonomic groups and geographical areas representing the vast diversity observed in squamates (see Figure 1). We also ran additional analyses using only those pairs for which the median of the age since divergence was equal to or lower than 10 million years ago (124 of snakes and 243 of lizards), a more conservative way to include species pairs with respect to potential loss of history to extinction.

**Figure 1.**
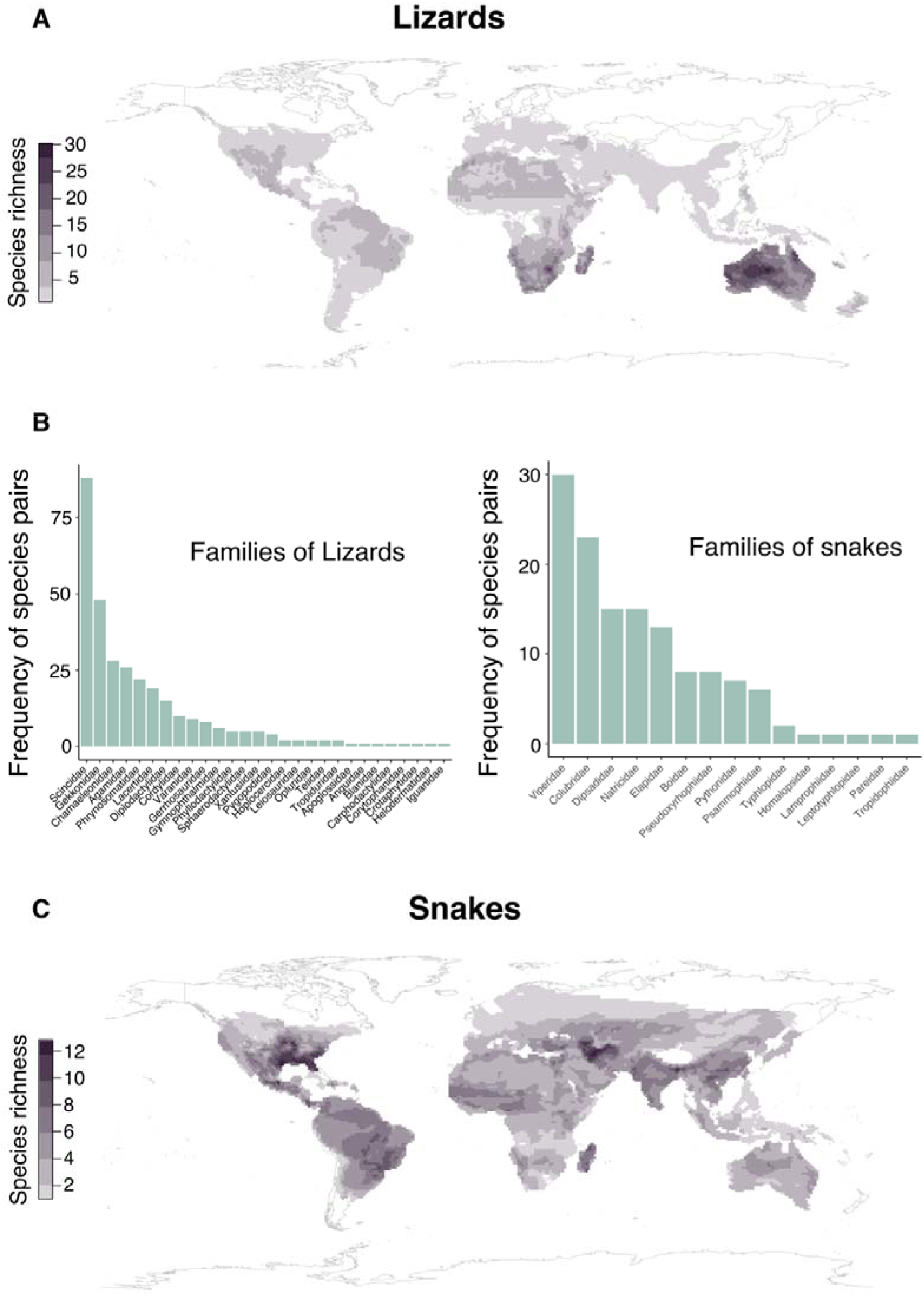
Species diversity analyzed in the present study across geographical regions (a, c) and the taxonomic diversity of the species pairs (b) of lizards and snakes included in our analyses.

### Geographical overlap

We considered a species pair as sympatric or allopatric based on the spatial overlap of their geographical ranges. We considered a pair as sympatric when more than 30% of the smaller species range overlapped with the range of the other species. We also ran additional analyses considering a given pair as sympatric when the smaller range overlapped more than 70%. These additional analyses allowed us to explore if our results on the coexistence dynamics of squamates would differ depending on how we define sympatry. We quantified geographic overlap using the Raster package in R (Hijmans, 2019). We should note that we choose a simpler geographical categorization for methodological purposes (see Statistical analyses), considering as “allopatric” species pairs that might have been originated under processes such as vicariance, parapatric speciation, or founder events (see Skeels & Cardillo, 2019).

### Geographical and ecological traits

#### Size of the geographical area

To explore if the size of the geographical area affects the tendency by which closely related species coexist over time, we categorized each species pair as occurring on islands or continents. We considered as islands all geographically isolated land masses smaller than Madagascar (the largest island considered). When the species comprising a given pair occurred both on islands and continents, we considered the geographical area as the one where both species occur. We did not have any cases where both the species occurred on islands and continents.

#### Ecological similarity

To quantify how ecologically similar two species are and, thus, to explore the importance of species interactions in affecting the tendency of closely related species to coexist over time, we measured specimens deposited in several scientific collections between the years of 2017 and 2019 (see supplementary methods). This massive data collection allowed us to obtain measurements for more than 2000 specimens of lizards and snakes comprising 300 and 152 species, respectively (150 and 76 sister pairs of lizards and snakes; Appendix II and III). Therefore, we were able to obtain morphological data for 47% and 57% of the pairs from our total pool (lizards and snakes, respectively). On average, we were able to measure 6 specimens per species of lizards and 5.7 specimens per species of snakes. For additional details on how we determined the sex and the sexual maturity of the specimens see the supplementary methods.

We focused on measuring morphological traits known to be associated with different axes of the ecological niches of squamates. Additionally, we aimed to include only males in our dataset as a way to decrease any noise caused by morphological differences resulting from sexual dimorphism. Measurements for lizards comprised snout-vent length (SVL), head length, width and height, jaw length, mid-body circumference, the length of the humerous, ulna, femur and tibia, and the distance between fore and hind limbs. Measurements for snakes comprised snout-vent length, head length, width and height, mid-body circumference, tail length and eye diameter. This dataset was combined with the morphological dataset for snakes of the family Viperidae generated by Alencar et al. (2017), and with a few measurements for snout-vent length taken from the literature (see Appendix II and III), all measurements from adult males only.

To explore the role of species interactions in driving the tendency of species to geographically overlap over time, we used two different metrics as proxies for ecological similarity of a given species pair. First, we calculated the absolute difference between the average of the log-transformed SVL of the species comprising each pair. The larger the difference between the SVL, the less ecologically similar species are expected to be (Wilson 1975; Ricklefs & Miles, 1994). We chose to use SVL as our metric for body size given it is traditionally used by herpetologists as a proxy for size (see Meiri, 2008). As a next step, we generated one morphospace for snakes and another for lizards but excluding legless lizards, and a second one for all lizards including legless lizards but excluding the distance between the fore and hind limbs measurement. To generate these morphospaces we first size-corrected the morphological variables (log-shape ratios) for each species by dividing each variable by the geometric mean calculated for each species (see below) and log-transforming the resulting value (Price et al., 2019; Friedman et al., 2020). The geometric mean was calculated for lizards as the cube root of the product of species average SVL, mid-body circumference and length of the limbs, and for snakes as the cube root of the product of species’ average SVL, tail length and mid-body circumference. We chose not to calculate the geometric mean across all morphological variables because most of them are inclusive measures of these dimensions of size (see Price et al., 2019). We chose to apply this size-correction method because it allows us to produce a metric that preserves the allometric component of shape (see Price et al., 2019). We performed phylogenetic principal component analyses using the R package phytools (Revell, 2009; Revell, 2012) to generate these morphospaces using the consensus phylogenetic tree from Tonini et al. (2016). Using these morphospaces, we calculated the Euclidean distance between species comprising each pair across all PhylPCA axes combined. The larger the euclidean distance, the less ecologically similar species are expected to be. In the end, we had two proxies for ecological similarity, (1) differences in SVL and (2) the Euclidean distance between species pairs within the morphospace.

To explore if the proxies for ecological similarity, that is, differences in SVL and differences in shape (Euclidean distance) were correlated, we performed PGLS analyses using ten phylogenetic trees from Tonini et al. (2016). These analyses suggested that despite being significantly correlated in lizards (R^2^ 0.59 – 0.77, p < 0.05), differences in the SVL and Euclidean distances are not correlated in snakes (R^2^ < 0.01, p > 0.1). In the subsequent analyses, we decided to use only the SVL differences for lizards and keep both the differences in SVL and the Euclidean distance as proxies for ecological similarity in snakes.

#### Dispersal ability

Despite often being considered a species trait, here we assess dispersal ability as a trait characterizing a given species pair due to methodological reasons (see statistical analyses below). To do this, we used two different metrics. First, we used the log-transformed ratio between the average range size of each pair and the age since the divergence of that pair. Range size can be a good proxy for dispersal ability because larger ranges might reflect a higher ability of lineages to geographically expand (Brown et al., 1996; Pigot et al., 2018). We decided to take into account the age since divergence between species comprising each pair because older lineages might have larger geographical ranges simply because they had more time to expand geographically and not as a result of dispersion ability. Therefore, the ratio between range size and age of divergence reflects the rate at which a given species pair was able to geographically expand. We calculated this ratio 100 times for each pair using the different age estimates obtained previously. As a next step, we used the average SVL for each pair. Because larger animals might also have larger home ranges and/or territories (Brown et al., 1996; Bonner, 2011), we expect that larger squamates might have higher dispersal ability. It is important to note that the analyses using average SVL as a proxy for dispersal ability included less species pairs compared to our first proxy given we measured a smaller number of species than our total species pool (see above).

To explore if the two proxies were correlated with each other, we performed PGLS analyses on the average snout-vent length and the average range size/age ratio, using ten randomly sampled sets of the latter and the ten corresponding phylogenetic trees from Tonini et al. (2016). Interestingly, all these analyses suggested that despite being significantly correlated (p < 0.05), the relationships have either very low R^2^ (0.06 – 0.08, for lizards) or moderate R^2^ (0.14 – 0.18, for snakes). For this reason, we decided to keep both proxies in our subsequent analyses (see below).

### Statistical analyses

We explored the coexistence dynamics of squamates over time by using different probabilistic models of species co-occurrence (Figure 2, Pigot et al., 2013, 2015). In this framework, the dynamic of coexistence is modelled as a constant rate Markov process, and maximum likelihood is used to perform model fitting and parameter estimates. In a general manner, the models allow us to calculate the probability that a species pair exists in its current geographical state (i.e. allopatry or sympatry) given their age since divergence and the parameters controlling the rates of transition from allopatry to sympatry (σ) and from sympatry to allopatry (ε) (Figure 2, see also Pigot & Tobias, 2015). Therefore, we were able to model how coexistence possibly changed over time by using a given set of species pairs. We fitted four different models to our dataset using the R package msm (Jackson, 2011): 1) allo-one-way, which assumes that lineages diverged in allopatry and then undergo a transition to sympatry (γ = 1, σ > 0, and ε = 0); 2) symp-one-way, which assumes that lineages diverged in sympatry and then undergo a transition to allopatry (γ = 0, σ = 0, and ε > 0); 3) allo-two-way, assumes that lineages diverged in allopatry, become sympatric but became allopatric again (γ = 1, σ > 0, ε > 0); Symp-two-way, which assumes that lineages diverged in sympatry, undergo a transition to allopatry but come back to sympatry later (γ = 0, σ > 0, and ε > 0).

**Figure 2.**
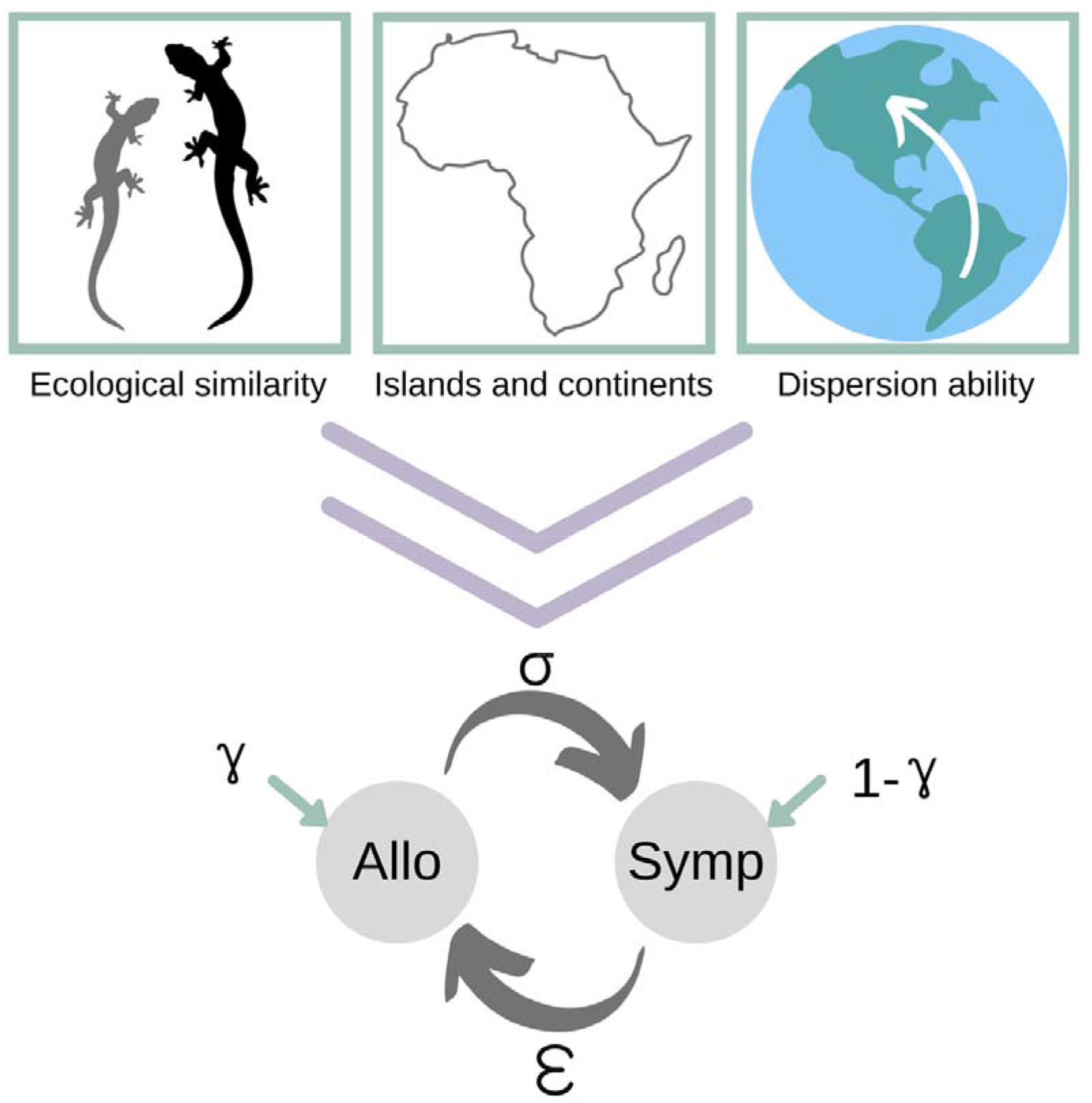
Summary of the statistical models compared in the present study and their parameters. γ = relative frequency of allopatric speciation, σ = transition rate from allopatry to sympatry, ε = transition rate from sympatry to allopatry. We included ecological similarity (i.e. body size and shape differences), occurrence on islands or continents, and the dispersion ability (i.e. range size/age ratio and mean body size) separately as covariates on the transition rate to sympatry. Modified from Pigot & Tobias (2015).

A schematic workflow of the model comparisons we performed is presented in Figure S1. First, we fitted all four models 100 times across species pairs of lizards and snakes separately using their age since divergence obtained from 100 different topologies (see above). We then evaluated how many times a given model was favored over the others using the Akaike Information Criterion (Burnham & Anderson, 2002) (see Table S1-S4). We considered the best model to be the one with the lowest AIC value and with a ΔAIC higher than two. To evaluate the effects of geographical position (island vs continent), ecological similarity (body size difference and euclidean distance) and dispersal ability (ratio of range area and age, average body size) in shaping the coexistence dynamics of squamates, we ran another set of analyses comparing all four models but also including the allo-one-way and the allo-two-way models with each candidate factor as a covariate of σ (the parameter describing the transition rate to sympatry). We decided not to include the Symp-two-way model with candidate factors as covariates in our model comparison because transitions to coexistence occur at a distinct stage in the symp-two-way model relative to those in the allopatric models. In summary, we performed model comparisons across the six models (allo-one-way, symp-one-way, allo-two-way, symp-two-way, allo-one-way with covariate, allo-two-way with covariate) five times for snakes and four times for lizards, including one distinct candidate driver in each of them (Table S1-S4, see summary in Figure S1). We performed model comparisons separately using each candidate factor as they frequently involved distinct species pairs datasets (e.g. morphology based proxies comprise fewer species pairs than range size, for example).

When candidate drivers significantly improved a model over the others, we quantified the hazard ratio to evaluate the direction of the relationship between transition rates to sympatry and the driver of coexistence (Table S5). To illustrate how transition rates to sympatry are affected by the drivers of coexistence, we extracted estimates of transition rates to sympatry under the best models selected and the corresponding hazard ratios for different values of the drivers (Figure 3). We performed posterior predictive simulations to evaluate how well the best models could actually predict the empirical data (see supplementary methods).

**Figure 3.**
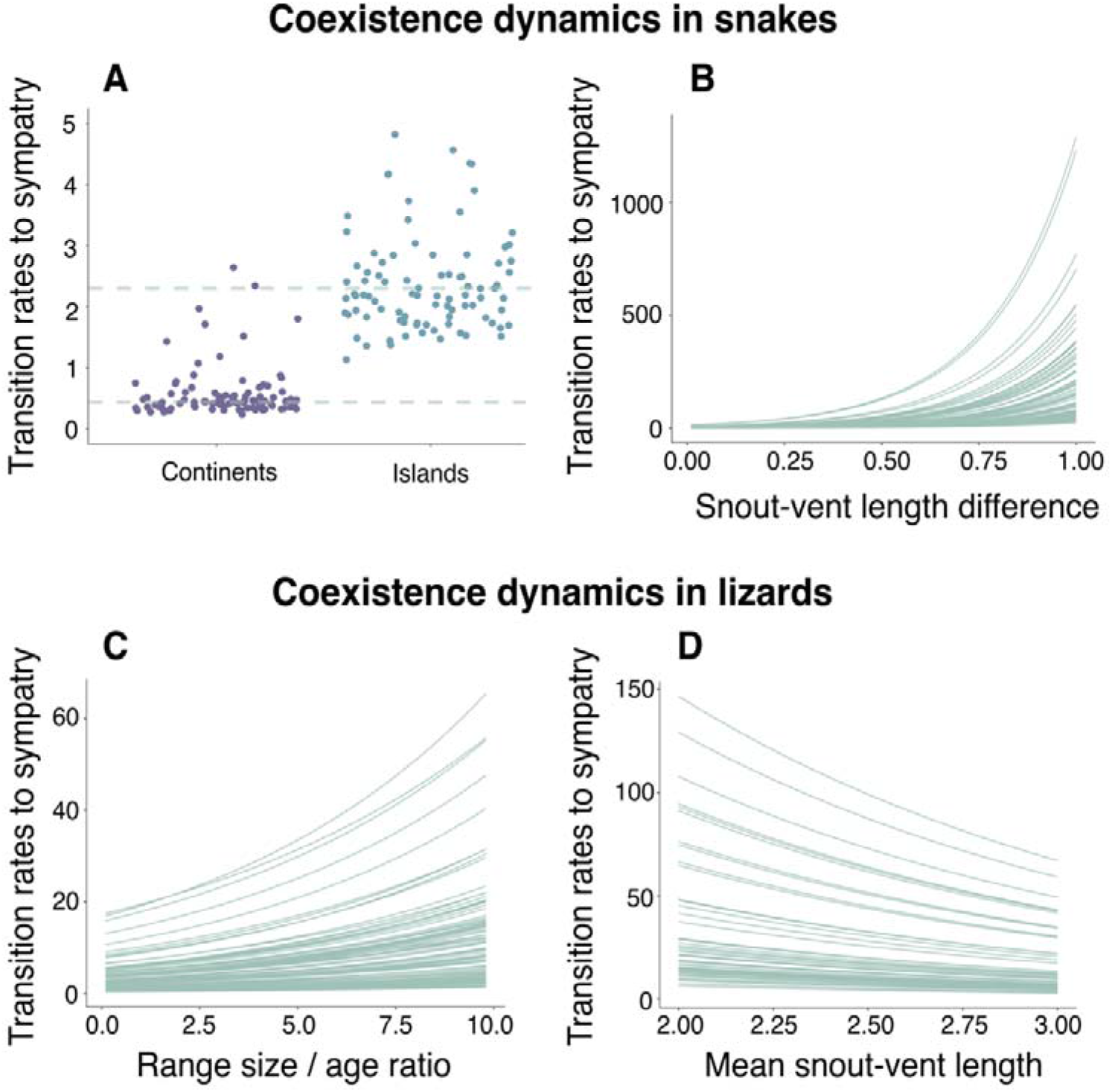
Estimates of transitions rates to sympatry relative to (a) species pairs occurrence on islands or continents, (b) snout-vent length difference, and (c) dispersal ability (range size / age ratio and mean snout-vent length), under the best model selected for snakes (a, b) and lizards (c, d). Multiple dots and lines represent estimates when using distinct age datasets (see Methods). Dashed lines in panel (a) represent medians of transition rates taken across all estimates. Given the extremely high values of 12 of these estimates (1 continent and 11 island), we excluded these from the plot for visualization purposes.

## Results

For our main dataset that considered sympatric species pairs as those with at least 30% of the smallest distribution overlapping and included pairs that diverged 20 million years ago or less, 37% and 48% of the species pairs are sympatric in snakes and lizards, respectively (Table S6). Proportions were very similar when considering only the species pairs with a median age since divergence of less than 10 mya (see Table S6). However, as expected, the number of sympatric pairs decreased when we considered as sympatric only those pairs with more than 70% of the smallest distribution overlapping (Table S6). Regardless of the dataset, however, lizards consistently show a higher proportion of sympatric pairs compared to snakes (Table S6). In general, sympatric pairs tend to be older than allopatric pairs among snakes (Table S7). In lizards, however, sympatric and allopatric species pairs have very similar ages of divergence, with sympatric pairs still being slightly older across datasets comprising pairs that have diverged 20 mya or less (see Table S7).

### Coexistence dynamics in squamates

#### Model selection without covariates

When comparing only the four models without adding any candidate traits that could potentially be driving coexistence dynamics, the “allo-two-way-model” was considered the best model for about 75% of the age datasets analyzed for snakes either when considering species pairs that diverged less than 20 or 10 mya (Table S1-S2). These results suggest that snakes generate species mainly via allopatric speciation with transitions to sympatry over time and back to allopatry as lineages become older. On the other hand, by comparing the four models without adding the covariates, we could not recover a unique best model for lizards (Figure S1-S2). The symp-two-way-model and the allo-two-way-model are equally likely, which suggests that geographical overlap in lizards changed considerably through time. Additionally, no best model was detected when performing model comparison with an overlap cut-off of 70% either in lizards or snakes (see Table S3-S4).

#### Model selection with the geographical setting as covariate

Being on an island or continent seems to be an important driver of species coexistence in snakes but not necessarily in lizards. When considering the two additional models that include the geographical setting as a covariate on the transition rate to sympatry, model comparisons suggested that the allo-two-way model with the covariate was the best model across all age datasets in snakes, regardless of only including younger pairs or not (Table S1-S2). According to this model, transition rates to sympatry substantially increase in species pairs of snakes occurring on islands compared to those occurring on continents (Figure 3, Table S5). When performing model comparison under the 70% overlap cut-off, the allo-two-way model with the covariate was still recovered as the best model across 76% of the age datasets in snakes. However, no single best model was recovered when considering only the younger pairs (Table S3-S4), although the model with geographical setting as covariate was still among the best models (it was tied with another model in 99 out of 100 comparisons). In contrast, no best model was selected for lizards and, therefore, we do not have evidence that being on an island or continent is relevant in driving their coexistence dynamics (Table S1-S4). It is also worth mentioning that, in general, the model that includes geographical setting as a covariate was frequently not among the best tied models in lizards, with the exception of when considering the 70% overlap threshold and species pairs that diverged less than 10 mya (33, 26, 56, or 97 out of 100 model comparisons depending on the dataset, see Tables S1-S4).

#### Model selection with ecological similarity as covariate

Ecological similarity seems to be an important driver of species coexistence in snakes but not in lizards. Model selection suggested that the allo-two-way model with body size difference as the covariate was the best model across all age datasets for snakes (all 100 model comparisons), regardless of whether the data were restricted to younger pairs (Table S1-S2). According to this model, transition rates to sympatry increase with increasing differences in body size between species pairs (Figure 3, Table S5). Despite always suggesting the same positive relationship, transition rates to sympatry and the hazard ratio have unreliably high values when estimated using ages of divergence extracted from some phylogenies (Figure 3, Table S5). This suggests that model fitting might have failed to converge in these datasets. However, simulations using parameter estimates under the best model are still able to recover proportions of sympatric and allopatric pairs similar to the empirical data (see below). When analyzing species pairs of snakes under the 70% overlap cut-off no model was preferred, but the model with body size difference as the covariate tied with another model about 60% of the time (Table S3-S4). In contrast, we did not recover a best model when adding the differences in shape (measured by the Euclidean distance, see Methods) of species pairs as the covariate on transition rates to sympatry in snakes, although this model is frequently tied with other models (see Tables S1-S4). Additionally, we did not find strong evidence of ecological similarity being an important driver of coexistence in lizards (Table S1-S4). It is important to mention, however, that although we did not recover a best model for lizards, the model with body size difference as the covariate is among the best models in several of the model comparisons performed (Table S1-S4).

#### Model selection with dispersal ability as covariate

Contrary to the analyses above, dispersal ability seems to be a relevant driver of coexistence dynamics in lizards but not snakes (Table S1-S2). Model selection suggested that the allo-two-way model with the ratio of range size and age as the covariate was the best model across more than 80% of the age datasets of lizards when considering the overlap threshold of 30%, regardless of whether only younger pairs were included (Table S1-S2). According to this model, transition rates to sympatry increase with faster rates of geographic expansion (i.e. the larger the ratio between geographical range and time; Figure 3, Table S5). We did not recover a single best model when analyzing species pairs of lizards under the 70% overlap cut-off (Table S3-S4), but the model with dispersal as a covariate was usually among the tied models (above 80% of the comparisons). When performing model selection with mean body size (the other proxy for dispersion ability) as the covariate, we recovered the allo-two-way model that included the covariate as the best model for 57% of the age datasets but only in our main analyses (30% overlap and all species pairs, Table S1). Surprisingly, however, according to this model transition rates to sympatry increase the smaller the species (Figure 3, Table S5). We did not find a single best model for these model comparisons using species pairs of snakes but models with dispersal proxies as covariates usually tied with other models (Table S1-S4).

### Posterior predictive simulations

For all best models with covariates selected in our main analyses (30% overlap between species and including all species pairs), simulated proportions of sympatric pairs frequently recovered the empirical proportions (Figure S2).

## Discussion

In lizards and snakes, neither time or sympatric speciation alone can explain the coexistence patterns observed among extant species. On the other hand, by performing extensive data collection and comparing statistical models, we found evidence for completely distinct drivers underlying the coexistence dynamics of these two closely related groups. In general, our results suggest that the species’ geographical setting, ecologically similarity, and dispersal ability might play different roles in shaping the coexistence dynamics within each taxonomic group analyzed.

By taking a first look at the patterns of geographical overlap, we found that most species pairs of snakes are allopatric. Allopatric speciation is indeed the rule across vertebrates, with many vertebrate groups having more allopatric than sympatric sister species (Barraclough & Vogler, 2000; Pigot & Tobias, 2015; García-Navas et al., 2020; see review in Hernández-Hernández et al., 2021). In lizards, however, allopatric and sympatric pairs had similar proportions. Allopatric pairs of snakes tend to be younger than sympatric ones, which conforms to a scenario where coexistence would be achieved with time. Indeed, without adding any covariates, the coexistence dynamics in snakes are best described by allopatric speciation with lineages eventually becoming sympatric as time goes by and becoming allopatric again as they get older (allo-two-way model, Table S1-S2). In lizards, on the other hand, sympatric species and allopatric pairs tend to have very similar ages, with allopatric pairs being older in some cases (Table S7). This result might suggest two scenarios: (1) sympatric speciation plays an important role in the diversification of lizards; (2) allopatric speciation is still the rule, but time alone does not explain the coexistence dynamics of lizards. In any case, we indeed could not recover a single best model in the first set of model comparisons for the group (Table S1-S4). It is worth noting that when a single model is chosen in lizards, the sympatric model is the one selected (19 vs 1, Table S1), and when two models are equally likely (80% of the time), the sympatric model is also among them (Table S1).

Taken together, these first results suggest that lizards might be more heterogeneous and have more dynamic distributions compared to snakes, either because their lineages are influenced by different processes and/or because their distributions change more through time. A wider array of processes acting simultaneously could potentially mask coexistence signals when we include all species in the same category “Lizards”. Indeed, lizards occurring in the same islands can show completely distinct patterns of diversification. At the Socotra Archipelago, for example, closely related species of the genera *Hemidactylus* and *Haemodracon* show sympatric distributions with marked differences in body size, whereas those of the genus *Pristurus* tend to be allopatric or parapatric with no apparent divergence in body size (see García-Porta et al., 2016). Furthermore, species richness can indeed be achieved via distinct pathways in distinct lizard clades and geographic regions. High species richness is associated with low functional divergence in agamids and gekkoes but with high functional diversity in varanids and scincids (Skeels et al., 2020). Exploring whether distinct clades are also characterized by distinct coexistence dynamics might be an interesting next step. However, we would need a much higher number of phylogenetically well-supported species pairs within each of the distinct clades to explore group-specific coexistence dynamics.

When adding covariates to the transition rates to sympatry, the allo-two-way model is frequently recovered as the best model across the analyses for which we were able to recover a single best model. In contrast to previous findings in birds (Pigot et al., 2018), islands seem to promote species coexistence in snakes. Surprisingly, however, this effect was not evident in lizards. In other words, the model with the geographical setting as covariate does not have higher support compared to those without the covariate in lizards. Indeed, in this set of analyses for lizards, ties were mainly between the allo-two-way and symp-two-way models that lack covariates (Table S1). In general, snakes have a slower life history and lower energetic demands compared to lizards (Pough, 1973; Vitt & Caldwell, 2014). Therefore, resources on islands might appear less scarce for snakes than lizards, potentially allowing snakes to more easily coexist in insular environments given the relaxation of some biotic constraints (e.g. predation, and perhaps less intense competition) and the smaller geographical limits of islands compared with continental settings. Lizards, on the other hand, have high population densities on islands (Novosolov et al., 2015) and the resultant scarcity of resources might lead to much more intense competition. Hence, the smaller geographical limits would not increase coexistence in insular lizards at least when compared with continental settings.

Species interactions also underlie species coexistence in snakes, and divergence in body size (but not shape) seems to be the most likely route to avoid competition and promote coexistence, similar to what has been suggested for birds (Pigot et al., 2018). On the other hand, we found no definitive or clear evidence that ecological similarity is a relevant driver of species coexistence in lizards. For 99% of the time when we had more than one best model (a tie) the model involving ecological divergence (complex-allo-two-way) was among them for lizards (Table S1). The prediction that the likelihood of coexistence increases with increasing the ecological difference between species, assumes that some species pairs would accumulate more morphological differences allopatrically allowing them to coexist faster than others (i.e. species sorting, Grant, 1972; Davies et al., 2007). Although we did not find definitive support for this prediction in lizards, morphological divergence could still be driving the coexistence dynamics at the local scale. If this is the case, we would expect that sympatric populations of a given species pair would be more morphologically distinct compared to allopatric populations of these same species (character displacement, Brown & Wilson, 1956; Davies et al., 2007). We might not have been able to capture these smaller-scale differences given our large-scale approach and goals. A fruitful next step would be to apply recently developed and promising frameworks to detangle the role of species sorting and character displacement in speciation and species coexistence dynamics (see Anderson & Weir, 2021), and to increase the intra-specific morphological sampling allowing us to compare morphological divergence at the population-level.

Even though competition has widely been seen as an important factor shaping community structure (e.g. Darwin, 1859; Webb et al., 2002; Cavender-Bares et al., 2009), a lack of strong evidence of competitive interactions shaping trait-divergence has also been suggested (e.g. Meiri et al., 2011; Stuart & Losos, 2013; Shi et al., 2018, Slavenko et al., 2021). Slavenko et al. (2021) suggested that environmental filtering might be more important in driving morphological divergence in lizards than competitive interactions, which could potentially explain why we did not find clear evidence for morphological differences mediating coexistence dynamics in lizards. Another possibility is that morphological divergence could be more relevant in some geographical regions than others or within some groups of lizards (see Skeels et al., 2020). Competition might be more prevalent in climatically stable areas (Dobzhansky, 1950; MacArthur, 1969; Henriques-Silva et al., 2019) or regions with lower resource availability and, therefore, morphological divergence could be an important driver of coexistence at lower compared with higher latitudes or on islands compared with continents (e.g. García-Porta et al., 2016), respectively.

In contrast to what we found for competitive interactions, dispersal ability (measured by the ratio between range size and age of divergence) seems to be a relevant driver of species coexistence in lizards but not necessarily in snakes. It is worth mentioning that our dispersal metric that uses range size/age has its limitations and assumptions. For example, it is thought that species might both start and go extinct with smaller range sizes (Foote, 2007). Hence we might be pulling together species at very different stages which might mask the potential effect of dispersal and perhaps explain the lack of dispersal effects in snakes. By restricting the time window of analysis, this problem might be ameliorated but not explicitly taken into account. However, the lack of an association is still present in snakes when considering species pairs that diverged less than 10 Mya.

Lizards indeed have been suggested to have comparatively higher dispersal abilities than snakes, at least regarding long-distance dispersal to oceanic islands (see Pitta et al., 2013). It is important to note, however, that we are not able to disentangle between the ability to move across geographical space and the ability to establish populations in new geographical areas (see Jønsson et al., 2016), as we considered both as being part of the “dispersal ability” of species. Simply being more mobile across space would not necessarily mean that species would successfully establish populations in new geographical areas, possibly allowing species coexistence. This scenario has the potential to be especially true for snakes, for which our results suggest that competitive interactions are relevant drivers of species coexistence. Therefore, in snakes, only those species that are different enough might be able to rapidly coexist, even if others have high dispersal abilities. The establishment of populations, and consequently coexistence among species, might also be compromised if not enough time has elapsed for strong reproductive barriers to emerge and prevent populations from fusing (Weir & Price, 2011). The interaction between dispersal and competition, as well as between competition and geographical setting, in driving species coexistence deserves further investigation (see Lowe & McPeek, 2014; Jønsson et al., 2016). Indeed, the statistical framework to explore the interaction between coexistence drivers is already available (Pigot et al., 2016, 2018). However, we would need a much higher number of phylogenetically well-supported species pairs, as well as their morphological information to be able to properly investigate these questions.

Despite being favored in some of the model comparisons, models with mean body size as the covariate on the transition rates to sympatry suggested that the smaller the lizard the higher these transition rates. This might look unexpected given that both proxies for dispersal ability (ratio between range and age, and body size) were positively, despite weakly, correlated (see Methods). However, the body size dataset comprises a much smaller number of sister pairs compared to the range and age ratio dataset, which could explain the differences in the coexistence dynamics depicted by the models.

On the other hand, we cannot rule out the possibility that the preconception that larger species would be better dispersers might not be totally true for lizards. Going further, the reason for a negative relationship between body size and the transition rates to sympatry could be interpreted in light of the well-known link between body size and specific axes of the ecological niche in animals (Peters, 1986; Meiri, 2008; Clarke, 2021). Contrary to what has been suggested for several other organisms, Costa et al. (2008) found that body size is negatively related to dietary niche-breadth in predatory lizards and, therefore, the diversity of prey consumed decreases as body size increases. Furthermore, several small lizard species are insectivorous (Pianka & Vitt, 2003) and might face less severe pressures when dealing with the scarcity of food resources compared to larger species that may rely on more limited food availability. This, in turn, might enable smaller species to share the same resources and coexist. Although some large bodied species of lizards are herbivorous, meaning that their food might be readily available in some environments, the higher total metabolic rate of larger animals also requires a greater caloric intake (Pough, 1973). Coupled with that, small insectivorous lizards can be more mobile than larger herbivorous due to the higher energetic food taken by the first (Pough, 1973). All of these factors could help to explain a higher incidence of coexistence over time in smaller lizards.

Going further, the relative importance of each coexistence driver in distinct stages of the coexistence process is an aspect that deserves to be explored in squamates. As found by Pigot et al. (2018), patterns of narrow geographic overlap in birds seem to be driven by dispersal abilities and age, whereas wider coexistence patterns are mainly driven by ecosystem productivity and niche-divergence. For several of the analyses performed here, especially when considering as sympatric the species pairs with a geographic overlap higher than 70% of the smallest distribution, model comparisons were not able to choose between models. This probably occurs because using the 70% cut-off inevitably decreases the number of sympatric pairs, probably affecting the ability of these analyses to discern between models. Therefore, the addition of more well-supported species pairs coupled with their morphological information is essential to detangle between the differential drivers of species coexistence in distinct moments in the past.

## Conclusions

This study shows that lizards and snakes, although closely related, differ greatly in the drivers of species coexistence. Speciation seems to predominantly occur via allopatric speciation in snakes. Lizards, on the other hand, seem to be more heterogeneous and have more dynamic distributions, which likely prevented us from recovering a single best model without the addition of covariates (see Results). In snakes, species that occur on islands or have different body sizes are more likely to coexist. In contrast, lizard species are more likely to co-occur when they have higher dispersal abilities.

Indeed, it has been widely shown that lizards tend to exhibit unique diversity patterns that frequently do not follow the “rules” that usually apply to snakes or other vertebrate groups (Meiri, 2007; Powney et al., 2010; Roll et al., 2017; Novosolov et al., 2018). These differences might have profound consequences either for community structure and lineage diversification, and care should be taken when analyzing these two taxonomic groups together (see also Slavenko et al., 2019; Whiting & Fox, 2020). Beyond this, our results emphasize that when analyzing biogeographical, macroecological or macroevolutionary patterns and processes, one should take into account not only the geographical scenario but also who these organisms are (see also White, 2016; Skeels et al., 2020). It is the interaction between where and who that will ultimately shape biodiversity patterns.

## Acknowledgements

We thank all the curators and staff from the scientific collections visited: K. De Queiroz, R. Wilson and A. Wynn (SNMNH), L. Scheinberg (CAS), F. Burbrink and D. Kizirian (AMNH), H. Zaher and A. Benetti (MZUSP), D. Rabosky and G. Schneider (UMMZ), J. Losos and J. Rosado (MCZ), J. McGuire and C. Spencer (MVZ), P. Doughty (WAM), J. Rowley, S. Mahony, C. Portway, T. Cutajar (AM). We also thank Lab MeMe members, especially G. Burin and D. Caetano, for their suggestions during the development of this study. We thank A. Pigot for kindly providing R codes and J.R. Hodge for proofreading. LRVA and TBQ thank Sao Paulo Research Foundation (FAPESP) for grants #2016/14292-1 and 2018/05462-6. This study also received a Visiting Collections Fellowship awarded by the Australian Museum Research Institute to LRVA.

## Data availability

Data used in this study are part of the supplementary material or have been published elsewhere.

## Notes

**Conflict of interest statement** The authors declare no competing of interests.

### Competing Interest Statement

The authors have declared no competing interest.

