## Supplementary material for "Geographical and ecological drivers of coexistence dynamics in squamate reptiles"

Supplementary methods.………………...………….…………………………………………..…2

Figure S1.………………...……………………………………………………………………..…4

Table S1.……….……………...………………………………………………………….……….5

Table S2.……….……………...………………………………………………………………..…8

Table S3.……….……………...…………………………………………………………………11

Table S4.……….……………...…………………………………………………………………14

Table S5.……….……………...…………………………………………………………………17

Table S6.……….……………...…………………………………………………………...….…18

Table S7.……….……………...……………………………………………………………....…18

Figure S2.………………...……………………………………………………………...…….…19

Appendix I.……….……….….…...………………………………………….......………….......20

Appendix II……………………………………………………………………………………....20

Appendix III…...………………………………………………………………………………....20

Supplementary methods

*Additional information on morphological data collection*

We analyzed specimens deposited in nine collections: Museum of Zoology of the University of São Paulo, Museum of Zoology of the University of Michigan, Museum of Comparative Zoology, Smithsonian National Museum of Natural History, American Museum of Natural History, California Academy of Sciences, Museum of Vertebrate Zoology, Western Australian Museum and Australian Museum (Appendix II and III). When possible, we tried to measure at least five adult males per species. The sex of the specimens was determined by the presence of hemipenis (when the specimen was already dissected) or using more indirect evidence such as, for example, size and shape of the tail, presence of well-developed pores and coloration in the cases where chromatic dimorphism is known to occur (e.g. *Sceloporus*) (the last two specific to lizards). To determine the sexual maturity of specimens, that is, whether they were adults or not, we used different approaches. When possible, we used literature information regarding SVL of adults to guide our decision. When the specimen was already dissected, we also examined the presence of well-developed testis and deferent ducts, which suggest that specimens might have reached sexual maturity. Well-developed pores and head ornaments in males (lizards) can also be indicative of sexual maturity and were also taken into account. It is important to note that we are working within a large taxonomic scale and analyzing specimens deposited in several different institutions. Scientific collections have their specific rules, which can vary between them. Additionally, basic natural history information regarding important aspects of squamates are missing for several species. Therefore, it would be impossible to use a single approach to determine sex and sexual maturity for such a large and diverse dataset.

*Simulation tests*

We performed posterior predictive simulations to evaluate how well the best models could predict the empirical data. We simulated coexistence dynamics across the empirical ages of species pairs using the parameter estimates obtained from the best models selected when considering as sympatric those species pairs with at least 30% of the smallest distribution overlapping and with age of divergence less than 20 mya (our main analyses). We performed these posterior predictive simulations to evaluate how reliable our results are for the models that took into account (1) the size of the geographical area in snakes, (2) ecological similarity (body size difference) in snakes, and (3) dispersion ability (both proxies) in lizards (see Results). We ran 100 simulations using each age of divergence dataset (total of 1000 simulations) for each best model selected using the function sim.fittedmsm from the R package msm, which simulates a dataset from a Markov model fitted using the maximum likelihood estimates as parameters and the same observation times as in the original data (Jackson 2011). We then compared the simulated and empirical proportion of sympatric species pairs.

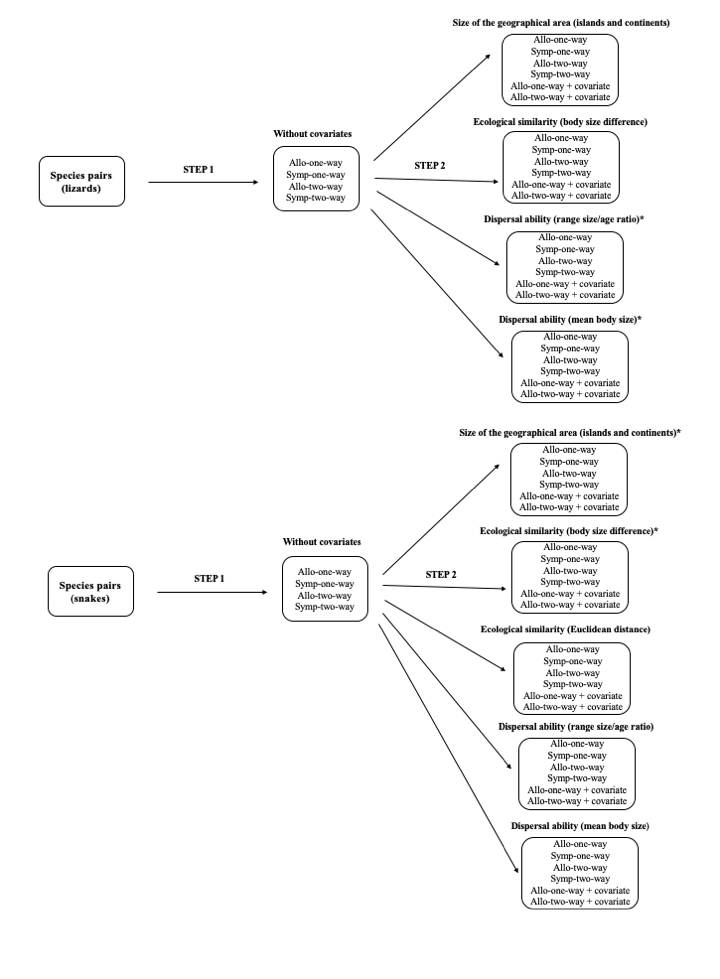

Figure S1. Schematic workflow of the model comparison approach. * indicates analyses for which the addition of drivers improved the allo-two-way model over all others.

Table S1. Results for model comparison across species pairs of snakes and lizards. In this set of analyses, our main one, we considered as sympatric the species pairs overlapping in more than 30% of the smallest distribution, and included those pairs that have diverged 20 million years ago or less (see Methods for details). “Simple model comparison” represents model comparison without including any of the factors as covariates on the transition rate to sympatry (σ). The other charts represent model comparisons performed by including one of the factors as covariates on the transition rates to sympatry. Complex-allo-one-way and Complex-allo-two-way represent the two models that included the factors as covariates. We considered to have a best model across the 100 model comparisons when the ∆AIC was higher than two in more than 50% of the age datasets analyzed (see Methods for details). For example, in the Simple model comparison (no covariates on σ), 19 model comparisons across the 100 age estimates of the species pairs suggested that the Symp-two-way model was the best one for lizards, whereas 76 suggested that the Allo-two-way model was the best one for snakes. Therefore, in this specific set of analyses, we could select the best model for snakes but not for lizards. Shaded area highlights when we considered to have a best model across model comparisons.

| Simple model comparison (no covariates on σ) | | | | |
| --- | --- | --- | --- | --- |
| **Lizards** | **Model** |  |  |  |
|  |  | **Best model** | **Not best model** | **Ties** |
|  | Simple Allo-one-way | 0 | 100 | 0 |
|  | Simple Symp-one-way | 0 | 100 | 0 |
|  | Simple Allo-two-way | 1 | 19 | 80 |
|  | Simple Symp-two-way | 19 | 1 | 80 |
| **Snakes** | **Model** |  |  |  |
|  |  | **Best model** | **Not best model** | **Ties** |
|  | Simple Allo-one-way | 0 | 100 | 0 |
|  | Simple Symp-one-way | 0 | 100 | 0 |
|  | Simple Allo-two-way | **76** | **0** | **24** |
|  | Simple Symp-two-way | 0 | 76 | 24 |
| Islands/continents as covariate on σ | | | | |
| **Lizards** | **Model** |  |  |  |
|  |  | **Best model** | **Not best model** | **Ties** |
|  | Simple Allo-one-way | 0 | 100 | 0 |
|  | Simple Symp-one-way | 0 | 100 | 0 |
|  | Simple Allo-two-way | 0 | 19 | 81 |
|  | Simple Symp-two-way | 19 | 1 | 80 |
|  | Complex-Allo-one-way | 0 | 100 | 0 |
|  | Complex-Allo-two-way | 0 | 67 | 33 |
| **Snakes** | **Model** |  |  |  |
|  |  | **Best model** | **Not best model** | **Ties** |
|  | Simple Allo-one-way | 0 | 100 | 0 |
|  | Simple Symp-one-way | 0 | 100 | 0 |
|  | Simple Allo-two-way | 0 | 100 | 0 |
|  | Simple Symp-two-way | 0 | 100 | 0 |
|  | Complex-Allo-one-way | 0 | 100 | 0 |
|  | **Complex-Allo-two-way** | **100** | **0** | **0** |
| Ecological similarity (body size difference) as covariate on σ | | | | |
| **Lizards** | **Model** |  |  |  |
|  |  | **Best model** | **Not best model** | **Ties** |
|  | Simple Allo-one-way | 0 | 100 | 0 |
|  | Simple Symp-one-way | 0 | 100 | 0 |
|  | Simple Allo-two-way | 0 | 7 | 93 |
|  | Simple Symp-two-way | 1 | 5 | 95 |
|  | Complex-Allo-one-way | 0 | 100 | 0 |
|  | Complex-Allo-two-way | 0 | 1 | 99 |
| **Snakes** | **Model** |  |  |  |
|  |  | **Best model** | **Not best model** | **Ties** |
|  | Simple Allo-one-way | 0 | 100 | 0 |
|  | Simple Symp-one-way | 0 | 100 | 0 |
|  | Simple Allo-two-way | 0 | 100 | 0 |
|  | Simple Symp-two-way | 0 | 100 | 0 |
|  | Complex-Allo-one-way | 0 | 100 | 0 |
|  | **Complex-Allo-two-way** | **100** | **0** | **0** |
| Ecological similarity (euclidean distance) as covariate on σ | | | | |
| **Snakes** | **Model** |  |  |  |
|  |  | **Best model** | **Not best model** | **Ties** |
|  | Simple Allo-one-way | 0 | 100 | 0 |
|  | Simple Symp-one-way | 0 | 100 | 0 |
|  | Simple Allo-two-way | 0 | 2 | 98 |
|  | Simple Symp-two-way | 2 | 1 | 97 |
|  | Complex-Allo-one-way | 0 | 100 | 0 |
|  | Complex-Allo-two-way | 0 | 2 | 98 |
| Dispersion ability (range size/age ratio) as covariate on σ | | | | |
| **Lizards** | **Model** |  |  |  |
|  |  | **Best model** | **Not best model** | **Ties** |
|  | Simple Allo-one-way | 0 | 100 | 0 |
|  | Simple Symp-one-way | 0 | 100 | 0 |
|  | Simple Allo-two-way | 0 | 100 | 0 |
|  | Simple Symp-two-way | 0 | 100 | 0 |
|  | Complex-Allo-one-way | 0 | 100 | 0 |
|  | **Complex-Allo-two-way** | **100** | **0** | **0** |
| **Snakes** | **Model** |  |  |  |
|  |  | **Best model** | **Not best model** | **Ties** |
|  | Simple Allo-one-way | 0 | 100 | 0 |
|  | Simple Symp-one-way | 0 | 100 | 0 |
|  | Simple Allo-two-way | 0 | 0 | 100 |
|  | Simple Symp-two-way | 0 | 76 | 24 |
|  | Complex-Allo-one-way | 0 | 100 | 0 |
|  | Complex-Allo-two-way | 0 | 6 | 94 |
| Dispersion ability (mean body size) as covariate on σ | | | | |
| **Lizards** | **Model** |  |  |  |
|  |  | **Best model** | **Not best model** | **Ties** |
|  | Simple Allo-one-way | 0 | 100 | 0 |
|  | Simple Symp-one-way | 0 | 100 | 0 |
|  | Simple Allo-two-way | 0 | 57 | 43 |
|  | Simple Symp-two-way | 0 | 100 | 0 |
|  | Complex-Allo-one-way | 0 | 100 | 0 |
|  | **Complex-Allo-two-way** | **57** | **0** | **43** |
| **Snakes** | **Model** |  |  |  |
|  |  | **Best model** | **Not best model** | **Ties** |
|  | Simple Allo-one-way | 0 | 100 | 0 |
|  | Simple Symp-one-way | 0 | 100 | 0 |
|  | Simple Allo-two-way | 0 | 2 | 98 |
|  | Simple Symp-two-way | 2 | 0 | 98 |
|  | Complex-Allo-one-way | 0 | 100 | 0 |
|  | Complex-Allo-two-way | 0 | 2 | 95 |

Table S2. Supplementary results for model comparison across species pairs of snakes and lizards considering as sympatric the species pairs overlapping in more than 30% of the smallest distribution but including only pairs that have diverged 10 million years ago or less. See legend of Table S1 and the Methods section for details.

| Simple model comparison (no covariates on σ) | | | | |
| --- | --- | --- | --- | --- |
| **Lizards** | **Model** |  |  |  |
|  |  | **Best model** | **Not best model** | **Ties** |
|  | Simple Allo-one-way | 0 | 100 | 0 |
|  | Simple Symp-one-way | 0 | 100 | 0 |
|  | Simple Allo-two-way | 2 | 32 | 66 |
|  | Simple Symp-two-way | 32 | 2 | 66 |
| **Snakes** | **Model** |  |  |  |
|  |  | **Best model** | **Not best model** | **Ties** |
|  | Simple Allo-one-way | 0 | 99 | 1 |
|  | Simple Symp-one-way | 0 | 100 | 0 |
|  | Simple Allo-two-way | **75** | **0** | **25** |
|  | Simple Symp-two-way | 0 | 76 | 24 |
| Island/continent as covariate on σ | | | | |
| **Lizards** | **Model** |  |  |  |
|  |  | **Best model** | **Not best model** | **Ties** |
|  | Simple Allo-one-way | 0 | 100 | 0 |
|  | Simple Symp-one-way | 0 | 100 | 0 |
|  | Simple Allo-two-way | 0 | 32 | 68 |
|  | Simple Symp-two-way | 32 | 2 | 66 |
|  | Complex-Allo-one-way | 0 | 100 | 0 |
|  | Complex-Allo-two-way | 0 | 74 | 26 |
| **Snakes** | **Model** |  |  |  |
|  |  | **Best model** | **Not best model** | **Ties** |
|  | Simple Allo-one-way | 0 | 100 | 0 |
|  | Simple Symp-one-way | 0 | 100 | 0 |
|  | Simple Allo-two-way | 0 | 100 | 0 |
|  | Simple Symp-two-way | 0 | 100 | 0 |
|  | Complex-Allo-one-way | 0 | 100 | 0 |
|  | **Complex-Allo-two-way** | **100** | **0** | **0** |
| Ecological similarity (body size difference) as covariate on σ | | | | |
| **Lizards** | **Model** |  |  |  |
|  |  | **Best model** | **Not best model** | **Ties** |
|  | Simple Allo-one-way | 0 | 100 | 0 |
|  | Simple Symp-one-way | 0 | 100 | 0 |
|  | Simple Allo-two-way | 0 | 20 | 80 |
|  | Simple Symp-two-way | 20 | 1 | 79 |
|  | Complex-Allo-one-way | 0 | 100 | 0 |
|  | Complex-Allo-two-way | 0 | 32 | 68 |
| **Snakes** | **Model** |  |  |  |
|  |  | **Best model** | **Not best model** | **Ties** |
|  | Simple Allo-one-way | 0 | 100 | 0 |
|  | Simple Symp-one-way | 0 | 100 | 0 |
|  | Simple Allo-two-way | 0 | 100 | 0 |
|  | Simple Symp-two-way | 0 | 100 | 0 |
|  | Complex-Allo-one-way | 0 | 100 | 0 |
|  | **Complex-Allo-two-way** | **100** | **0** | **0** |
| Ecological similarity (euclidean distance) as covariate on σ | | | | |
| **Snakes** | **Model** |  |  |  |
|  |  | **Best model** | **Not best model** | **Ties** |
|  | Simple Allo-one-way | 0 | 100 | 0 |
|  | Simple Symp-one-way | 0 | 100 | 0 |
|  | Simple Allo-two-way | 0 | 2 | 98 |
|  | Simple Symp-two-way | 1 | 5 | 94 |
|  | Complex-Allo-one-way | 0 | 100 | 0 |
|  | Complex-Allo-two-way | 0 | 1 | 99 |
| Dispersion ability (range size/age ratio) as covariate on σ | | | | |
| **Lizards** | **Model** |  |  |  |
|  |  | **Best model** | **Not best model** | **Ties** |
|  | Simple Allo-one-way | 0 | 100 | 0 |
|  | Simple Symp-one-way | 0 | 100 | 0 |
|  | Simple Allo-two-way | 0 | 100 | 0 |
|  | Simple Symp-two-way | 0 | 81 | 19 |
|  | Complex-Allo-one-way | 0 | 100 | 0 |
|  | **Complex-Allo-two-way** | **81** | **0** | **19** |
| **Snakes** | **Model** |  |  |  |
|  |  | **Best model** | **Not best model** | **Ties** |
|  | Simple Allo-one-way | 0 | 99 | 1 |
|  | Simple Symp-one-way | 0 | 100 | 0 |
|  | Simple Allo-two-way | 0 | 0 | 100 |
|  | Simple Symp-two-way | 0 | 76 | 24 |
|  | Complex-Allo-one-way | 0 | 100 | 0 |
|  | Complex-Allo-two-way | 0 | 4 | 96 |
| Dispersion ability (mean body size) as covariate on σ | | | | |
| **Lizards** | **Model** |  |  |  |
|  |  | **Best model** | **Not best model** | **Ties** |
|  | Simple Allo-one-way | 0 | 100 | 0 |
|  | Simple Symp-one-way | 0 | 100 | 0 |
|  | Simple Allo-two-way | 0 | 20 | 80 |
|  | Simple Symp-two-way | 20 | 1 | 79 |
|  | Complex-Allo-one-way | 0 | 100 | 0 |
|  | Complex-Allo-two-way | 0 | 21 | 79 |
| **Snakes** | **Model** |  |  |  |
|  |  | **Best model** | **Not best model** | **Ties** |
|  | Simple Allo-one-way | 0 | 100 | 0 |
|  | Simple Symp-one-way | 0 | 100 | 0 |
|  | Simple Allo-two-way | 0 | 2 | 98 |
|  | Simple Symp-two-way | 2 | 0 | 98 |
|  | Complex-Allo-one-way | 0 | 100 | 0 |
|  | Complex-Allo-two-way | 0 | 3 | 97 |

Table S3. Supplementary results for model comparison across species pairs of snakes and lizards considering as sympatric the species pairs overlapping in more than 70% of the smallest distribution and including pairs that have diverged 20 million years ago or less. See legend of Table S1 and the Methods section for details.

| Simple model comparison (no covariates on σ) | | | | |
| --- | --- | --- | --- | --- |
| **Lizards** | **Model** |  |  |  |
|  |  | **Best model** | **Not best model** | **Ties** |
|  | Simple Allo-one-way | 0 | 100 | 0 |
|  | Simple Symp-one-way | 0 | 100 | 0 |
|  | Simple Allo-two-way | 6 | 1 | 93 |
|  | Simple Symp-two-way | 1 | 6 | 93 |
| **Snakes** | **Model** |  |  |  |
|  |  | **Best model** | **Not best model** | **Ties** |
|  | Simple Allo-one-way | 0 | 99 | 1 |
|  | Simple Symp-one-way | 0 | 100 | 0 |
|  | Simple Allo-two-way | 16 | 0 | 84 |
|  | Simple Symp-two-way | 0 | 16 | 84 |
| Islands/continents as covariate on σ | | | | |
| **Lizards** | **Model** |  |  |  |
|  |  | **Best model** | **Not best model** | **Ties** |
|  | Simple Allo-one-way | 0 | 100 | 0 |
|  | Simple Symp-one-way | 0 | 100 | 0 |
|  | Simple Allo-two-way | 0 | 69 | 31 |
|  | Simple Symp-two-way | 0 | 74 | 26 |
|  | Complex-Allo-one-way | 0 | 100 | 0 |
|  | Complex-Allo-two-way | 44 | 0 | 56 |
| **Snakes** | **Model** |  |  |  |
|  |  | **Best model** | **Not best model** | **Ties** |
|  | Simple Allo-one-way | 0 | 100 | 0 |
|  | Simple Symp-one-way | 0 | 100 | 0 |
|  | Simple Allo-two-way | 0 | 80 | 20 |
|  | Simple Symp-two-way | 0 | 96 | 4 |
|  | Complex-Allo-one-way | 0 | 100 | 0 |
|  | **Complex-Allo-two-way** | **76** | **0** | **24** |
| Ecological similarity (body size difference) as covariate on σ | | | | |
| **Lizards** | **Model** |  |  |  |
|  |  | **Best model** | **Not best model** | **Ties** |
|  | Simple Allo-one-way | 0 | 100 | 0 |
|  | Simple Symp-one-way | 0 | 100 | 0 |
|  | Simple Allo-two-way | 0 | 0 | 100 |
|  | Simple Symp-two-way | 0 | 4 | 96 |
|  | Complex-Allo-one-way | 0 | 100 | 0 |
|  | Complex-Allo-two-way | 0 | 10 | 90 |
| **Snakes** | **Model** |  |  |  |
|  |  | **Best model** | **Not best model** | **Ties** |
|  | Simple Allo-one-way | 0 | 100 | 0 |
|  | Simple Symp-one-way | 0 | 100 | 0 |
|  | Simple Allo-two-way | 0 | 6 | 94 |
|  | Simple Symp-two-way | 6 | 0 | 94 |
|  | Complex-Allo-one-way | 0 | 100 | 0 |
|  | Complex-Allo-two-way | 0 | 38 | 62 |
| Ecological similarity (euclidean distance) as covariate on σ | | | | |
| **Snakes** | **Model** |  |  |  |
|  |  | **Best model** | **Not best model** | **Ties** |
|  | Simple Allo-one-way | 0 | 100 | 0 |
|  | Simple Symp-one-way | 0 | 100 | 0 |
|  | Simple Allo-two-way | 0 | 6 | 94 |
|  | Simple Symp-two-way | 6 | 0 | 100 |
|  | Complex-Allo-one-way | 0 | 100 | 0 |
|  | Complex-Allo-two-way | 0 | 14 | 86 |
| Dispersion ability (range size/age ratio) as covariate on σ | | | | |
| **Lizards** | **Model** |  |  |  |
|  |  | **Best model** | **Not best model** | **Ties** |
|  | Simple Allo-one-way | 0 | 100 | 0 |
|  | Simple Symp-one-way | 0 | 100 | 0 |
|  | Simple Allo-two-way | 0 | 1 | 99 |
|  | Simple Symp-two-way | 1 | 5 | 94 |
|  | Complex-Allo-one-way | 0 | 100 | 0 |
|  | Complex-Allo-two-way | 0 | 13 | 87 |
| **Snakes** | **Model** |  |  |  |
|  |  | **Best model** | **Not best model** | **Ties** |
|  | Simple Allo-one-way | 0 | 99 | 1 |
|  | Simple Symp-one-way | 0 | 100 | 0 |
|  | Simple Allo-two-way | 0 | 0 | 100 |
|  | Simple Symp-two-way | 0 | 16 | 84 |
|  | Complex-Allo-one-way | 0 | 99 | 1 |
|  | Complex-Allo-two-way | 0 | 2 | 98 |
| Dispersion ability (mean body size) as covariate on σ | | | | |
| **Lizards** | **Model** |  |  |  |
|  |  | **Best model** | **Not best model** | **Ties** |
|  | Simple Allo-one-way | 0 | 100 | 0 |
|  | Simple Symp-one-way | 0 | 100 | 0 |
|  | Simple Allo-two-way | 0 | 0 | 100 |
|  | Simple Symp-two-way | 1 | 35 | 64 |
|  | Complex-Allo-one-way | 0 | 100 | 0 |
|  | Complex-Allo-two-way | 0 | 0 | 100 |
| **Snakes** | **Model** |  |  |  |
|  |  | **Best model** | **Not best model** | **Ties** |
|  | Simple Allo-one-way | 0 | 100 | 0 |
|  | Simple Symp-one-way | 0 | 100 | 0 |
|  | Simple Allo-two-way | 0 | 6 | 94 |
|  | Simple Symp-two-way | 6 | 0 | 94 |
|  | Complex-Allo-one-way | 0 | 100 | 0 |
|  | Complex-Allo-two-way | 0 | 39 | 61 |

Table S4. Supplementary results for the model comparison across species pairs of snakes and lizards considering as sympatric the species pairs overlapping in more than 70% of the smallest distribution but including only pairs that have diverged 10 million years ago or less. See legend of Table S1 and the Methods section for details.

| Simple model comparison (no covariates on σ) | | | | |
| --- | --- | --- | --- | --- |
| **Lizards** | **Model** |  |  |  |
|  |  | **Best model** | **Not best model** | **Ties** |
|  | Simple Allo-one-way | 0 | 100 | 0 |
|  | Simple Symp-one-way | 0 | 100 | 0 |
|  | Simple Allo-two-way | 5 | 1 | 94 |
|  | Simple Symp-two-way | 1 | 5 | 94 |
| **Snakes** | **Model** |  |  |  |
|  |  | **Best model** | **Not best model** | **Ties** |
|  | Simple Allo-one-way | 0 | 98 | 2 |
|  | Simple Symp-one-way | 0 | 100 | 0 |
|  | Simple Allo-two-way | 16 | 0 | 84 |
|  | Simple Symp-two-way | 0 | 17 | 83 |
| Islands/continents as covariate on σ | | | | |
| **Lizards** | **Model** |  |  |  |
|  |  | **Best model** | **Not best model** | **Ties** |
|  | Simple Allo-one-way | 0 | 100 | 0 |
|  | Simple Symp-one-way | 0 | 100 | 0 |
|  | Simple Allo-two-way | 0 | 1 | 99 |
|  | Simple Symp-two-way | 1 | 5 | 94 |
|  | Complex-Allo-one-way | 0 | 100 | 0 |
|  | Complex-Allo-two-way | 0 | 0 | 97 |
| **Snakes** | **Model** |  |  |  |
|  |  | **Best model** | **Not best model** | **Ties** |
|  | Simple Allo-one-way | 0 | 98 | 2 |
|  | Simple Symp-one-way | 0 | 100 | 0 |
|  | Simple Allo-two-way | 0 | 0 | 100 |
|  | Simple Symp-two-way | 0 | 22 | 78 |
|  | Complex-Allo-one-way | 0 | 99 | 1 |
|  | Complex-Allo-two-way | 0 | 1 | 99 |
| Ecological similarity (body size difference) as covariate on σ | | | | |
| **Lizards** | **Model** |  |  |  |
|  |  | **Best model** | **Not best model** | **Ties** |
|  | Simple Allo-one-way | 0 | 100 | 0 |
|  | Simple Symp-one-way | 0 | 100 | 0 |
|  | Simple Allo-two-way | 0 | 0 | 100 |
|  | Simple Symp-two-way | 0 | 2 | 98 |
|  | Complex-Allo-one-way | 0 | 100 | 0 |
|  | Complex-Allo-two-way | 0 | 21 | 79 |
| **Snakes** | **Model** |  |  |  |
|  |  | **Best model** | **Not best model** | **Ties** |
|  | Simple Allo-one-way | 0 | 99 | 1 |
|  | Simple Symp-one-way | 0 | 100 | 0 |
|  | Simple Allo-two-way | 0 | 7 | 93 |
|  | Simple Symp-two-way | 7 | 0 | 100 |
|  | Complex-Allo-one-way | 0 | 100 | 0 |
|  | Complex-Allo-two-way | 0 | 39 | 61 |
| Ecological similarity (euclidean distance) as covariate on σ | | | | |
| **Snakes** | **Model** |  |  |  |
|  |  | **Best model** | **Not best model** | **Ties** |
|  | Simple Allo-one-way | 0 | 100 | 0 |
|  | Simple Symp-one-way | 0 | 100 | 0 |
|  | Simple Allo-two-way | 0 | 7 | 93 |
|  | Simple Symp-two-way | 7 | 0 | 93 |
|  | Complex-Allo-one-way | 0 | 100 | 0 |
|  | Complex-Allo-two-way | 0 | 32 | 68 |
| Dispersion ability (range size/age ratio) as covariate on σ | | | | |
| **Lizards** | **Model** |  |  |  |
|  |  | **Best model** | **Not best model** | **Ties** |
|  | Simple Allo-one-way | 0 | 100 | 0 |
|  | Simple Symp-one-way | 0 | 100 | 0 |
|  | Simple Allo-two-way | 0 | 1 | 99 |
|  | Simple Symp-two-way | 1 | 4 | 95 |
|  | Complex-Allo-one-way | 0 | 100 | 0 |
|  | Complex-Allo-two-way | 0 | 16 | 84 |
| **Snakes** | **Model** |  |  |  |
|  |  | **Best model** | **Not best model** | **Ties** |
|  | Simple Allo-one-way | 0 | 98 | 2 |
|  | Simple Symp-one-way | 0 | 100 | 0 |
|  | Simple Allo-two-way | 0 | 1 | 99 |
|  | Simple Symp-two-way | 0 | 17 | 83 |
|  | Complex-Allo-one-way | 0 | 97 | 3 |
|  | Complex-Allo-two-way | 0 | 2 | 98 |
| Dispersion ability (mean body size) as covariate on σ | | | | |
| **Lizards** | **Model** |  |  |  |
|  |  | **Best model** | **Not best model** | **Ties** |
|  | Simple Allo-one-way | 0 | 100 | 0 |
|  | Simple Symp-one-way | 0 | 100 | 0 |
|  | Simple Allo-two-way | 0 | 0 | 100 |
|  | Simple Symp-two-way | 0 | 2 | 98 |
|  | Complex-Allo-one-way | 0 | 100 | 0 |
|  | Complex-Allo-two-way | 0 | 1 | 99 |
| **Snakes** | **Model** |  |  |  |
|  |  | **Best model** | **Not best model** | **Ties** |
|  | Simple Allo-one-way | 0 | 99 | 1 |
|  | Simple Symp-one-way | 0 | 100 | 0 |
|  | Simple Allo-two-way | 0 | 7 | 93 |
|  | Simple Symp-two-way | 7 | 0 | 93 |
|  | Complex-Allo-one-way | 0 | 100 | 0 |
|  | Complex-Allo-two-way | 0 | 39 | 61 |

Table S5. Hazard ratios estimated under the best models across the 100 age of divergence datasets. Hazard ratios greater than or less than one indicate positive and negative effects on coexistence, respectively (Pigot et al. 2018). Numbers in parenthesis correspond to the lower and upper bound of 95% confidence limits estimated across the 100 age of divergence datasets analyzed. Values presented correspond to the mean calculated across the age of divergence datasets.

|  |  | **Hazard ratio** | |
| --- | --- | --- | --- |
| **30% overlap** | | **20 mya** | **10 mya** |
| Snakes | Island occurrence | 5.28 (2.04 - 13.70) | 4.10 (1.56 - 10.78) |
|  | Ecological similarity (body size diff) | 96.87 (2.74 - 3425.86) | 132.61 (2.92 - 6047.95) |
| Lizards | Dispersal ability (range size/age) | 1.14 (1.04 - 1.25) | 1.13 (1.03 - 1.25) |
|  | Dispersal ability (mean body size) | 0.46 (0.22 - 0.97) | - |
| **70% overlap** | | **20 mya** | **10 mya** |
| Snakes | Size of the geographical area | 2.73 (1.07 - 6.96) | - |

Table S6. Proportion of sympatric and allopatric species pairs across the different datasets we analyzed. These distinct datasets comprised the different ways we categorized species pairs as sympatric (30% or 70% of overlap) and including species pairs with medians of ages since divergence with a maximum of 20 mya (our main analyses) or 10 mya (see Methods for details). Mya: Million years ago.

| **Lizards** |  | 30% overlap | | 70% overlap | |
| --- | --- | --- | --- | --- | --- |
|  |  | 20 mya | 10mya | 20mya | 10mya |
|  | Sympatric | 48% | 46% | 31% | 30% |
|  | Allopatric | 52% | 54% | 69% | 70% |
| **Snakes** |  | 30% overlap | | 70% overlap | |
|  |  | 20 mya | 10mya | 20mya | 10mya |
|  | Sympatric | 37% | 37% | 24% | 24% |
|  | Allopatric | 63% | 63% | 76% | 76% |

Table S7. Median of the age since divergence across sympatric and allopatric species pairs for each dataset used in the present study. Medians were calculated by taking into account 100 age estimates for each species pair (see Methods for details). Mya: Million years ago.

| **Lizards** |  | 30% overlap | | 70% overlap | |
| --- | --- | --- | --- | --- | --- |
|  |  | 20 mya | 10mya | 20mya | 10mya |
|  | Sympatric | 5.84 | 4.15 | 5.92 | 4.27 |
|  | Allopatric | 5.80 | 4.75 | 5.82 | 4.57 |
| **Snakes** |  | 30% overlap | | 70% overlap | |
|  |  | 20 mya | 10mya | 20mya | 10mya |
|  | Sympatric | 4.40 | 4.15 | 4.40 | 4.11 |
|  | Allopatric | 3.43 | 3.14 | 3.70 | 3.37 |

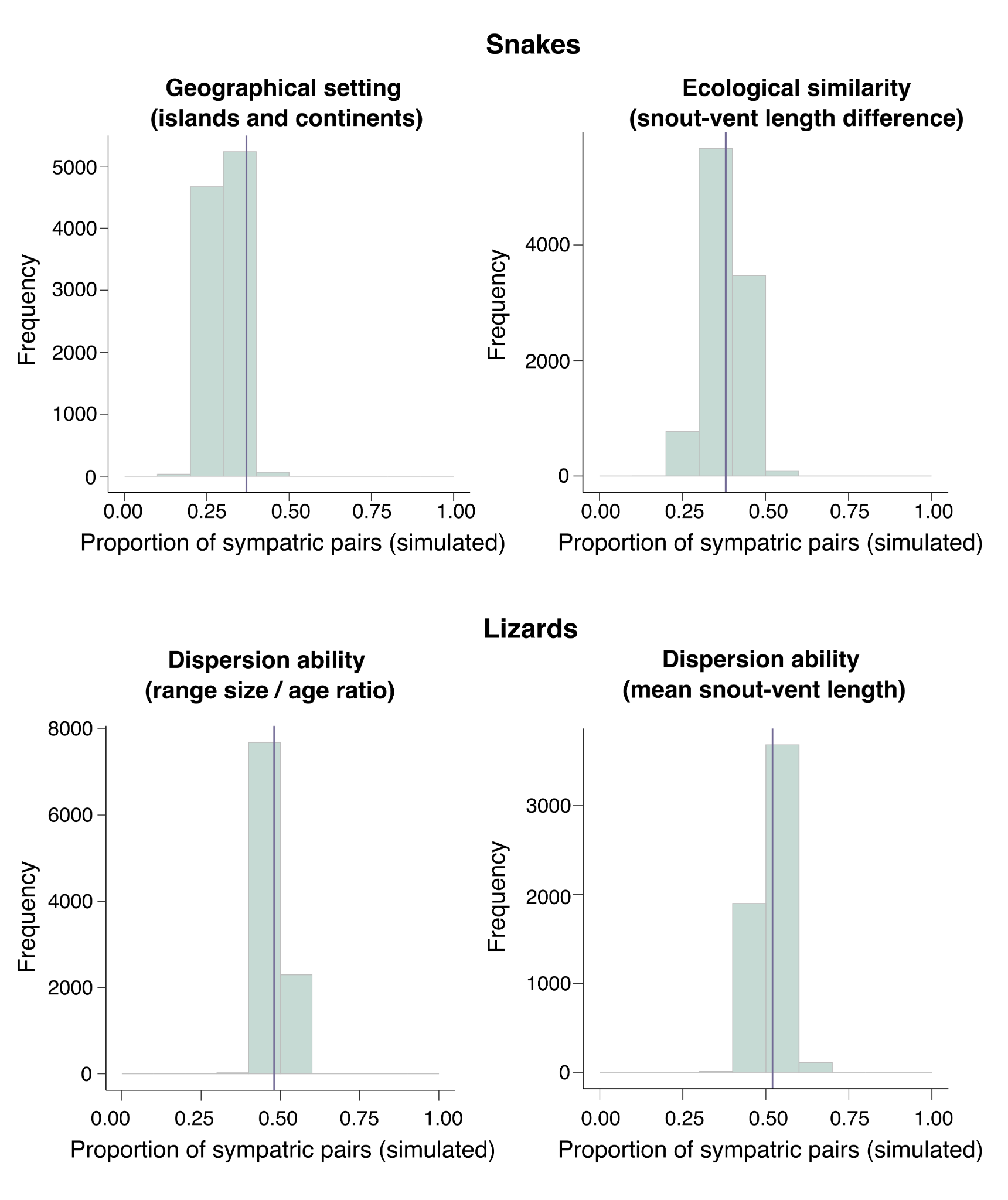

Figure S2. Results for the posterior predictive simulations performed using parameter estimates from the best model selected for snakes and lizards considering as sympatric the species pairs overlapping in more than 30% of the smallest distribution and including pairs that have diverged 20 million years ago or less (our main analyses). Vertical lines correspond to the empirical proportion of sympatric species.

**APPENDIX**

Appendix I – Excel file comprising all species pairs analyzed in the present study, their geographical overlap categorization using different sympatric thresholds (30% and 70%) and their occurrence on islands or continents.

Appendix II – Excel file with the list of all the specimens analyzed in this study with their corresponding catalog number.

Appendix III – Excel file with the morphological information and the number of specimens analyzed per species. SVL = snout-vent length, TL = tail length, HL = head length, HW = head width, HH = head height, ED = eye diameter, JL = Jaw length, CIRC = body circumference, HumL = length of the humerous, UlnL = length of the Ulna, FemL = length of the femur, TibL = length of the tibia, FHLDist = fore hind limb distance.
